# Unbiased whole-genome deep sequencing of human and porcine stool samples reveals circulation of multiple groups of rotaviruses and a putative zoonotic infection

**DOI:** 10.1101/058875

**Authors:** My VT Phan, Pham Hong Anh, Nguyen Van Cuong, Bas B. Oude Munnink, Lia van der Hoek, Phuc Tran My, Tue Ngo Tri, Juliet E. Bryant, Stephen Baker, Guy Thwaites, Mark Woolhouse, Paul Kellam, Maia A. Rabaa, Matthew Cotten on behalf of the VIZIONS Consortium, on behalf of the VIZIONS Consortium

**Affiliations:** Virus Genomics, Wellcome Trust Sanger Institute, Hinxton, Cambridge, United Kingdom; Oxford University Clinical Research Unit, Ho Chi Minh City, Vietnam; Laboratory of Experimental Virology, Academic Medical Center, University of Amsterdam, Amsterdam, the Netherlands; Centre for Tropical Medicine, Nuffield Department of Medicine, University of Oxford, Oxford, United Kingdom; London School of Tropical Medicine and Hygiene, London, United Kingdom; Centre for Immunity, Infection & Evolution, University of Edinburgh, Edinburgh, United Kingdom; Kymab Inc; Imperial College, London

**Author notes:** Corresponding authors: Paul Kellam and Matthew Cotten, Virus genomics, Wellcome Trust Sanger Institute, Hinxton Cambridge CB10 1SA, United Kingdom.

**Keywords:** rotavirus, deep sequencing, whole genomes, virus surveillance, zoonotic infection

## Abstract

Coordinated and synchronous virological surveillance for zoonotic viruses in both human clinical cases and animal reservoirs provides an opportunity to identify interspecies virus movement. Rotavirus is an important cause of viral gastroenteritis in humans and animals. We have documented the rotavirus diversity within co-located humans and animals sampled from the Mekong delta region of Vietnam using a primer-independent, agnostic, deep sequencing approach. A total of 296 stool samples (146 from diarrhoeal human patients and 150 from pigs living in the same geographical region) were directly sequenced, generating the genomic sequences of 60 human rotaviruses (all group A) and 31 porcine rotaviruses (13 group A, 7 group B, 6 group C and 5 group H). Phylogenetic analyses showed the co-circulation of multiple distinct rotavirus group A (RVA) genotypes/strains, many of which were divergent from the strain components of licensed RVA vaccines, as well as considerable virus diversity in pigs including full genomes of rotaviruses in groups B, C and H, none of which have been previously reported in Vietnam. Furthermore the detection of an atypical RVA genotype constellation (G4-P[6]-I1-R1-C1-M1-A8-N1-T7-E1-H1) in a human patient and a pig from the same region provides some evidence for a zoonotic event

## INTRODUCTION

Rotavirus (RV) infections are the leading cause of acute gastroenteritis globally, with a disproportionally greater morbidity and mortality in developing countries of Asia and sub-Saharan Africa (1, 2). RV can infect humans and different animal species and is considered, in part, a zoonotic disease in humans (3). Rotavirus zoonotic infections and transmissions, have been shown with animal strains moving into humans via direct contact with animals or exposure to environmental contamination (4–6), and present a challenge to infection control and management. RV is a non-enveloped double-stranded RNA virus forming the single genus *Rotavirus* in the Reoviridae family, with a 18.5 kb genome of 11 segments encoding six structural (VP1-4, VP6 and VP7) and five or six non-structural proteins (NSP1-NSP5/6) (3, 7). RVs are classified into eight established groups (A - H) and a new tentative group (I) based on the genetic and antigenic differences of VP6, with viruses of group A, B, C and H known to infect both humans and other animals (8, 9).

Within each rotavirus group, strains are distinguished and classified into G and P genotypes based on VP7 surface glycoprotein and VP4 spike protein, respectively (3). For rotavirus group A (RVA), at least 27 G and 37 P types have been detected in human and animals, with typical combinations of G-types (G1-G4, G9, G12) and P-types (P[4], P[6], P[8]) found in human infections globally, and different combinations of - and P-types (G3-G5, G9, G11, P[6], P[7], P[13]) commonly found in pigs (6, 10–13). With considerable more sequence data available for RVA as compared to non-RVA strains, a genotype classification system based on 11 genomic segments is recommended for RVA, providing genomic insights into genotype constellations of both common and novel human and animal RVA strains (14). In this RVA genotyping system, Gx-P[x]-Ix-Rx-Cx-Mx-Ax-Nx-Tx-Ex-Hx represents the genotypes of VP7-VP4-VP6-VP1-VP2-VP3-NSP1-NSP2-NSP3-NSP4-NSP5 segments, with the most prevalent human strains belonging to constellations of Wa-like (I1-R1-C1-M1-A1-N1-T1-E1-H1, commonly associated with G1P[8], G3P[8], G4P[8], G9P[8], G12P[8]) and DS-1-like (I2-R2-C2-M2-A2-N2-T2-E2-H2, commonly combined with G2P[4]) (14, 15). Using this system, a common origin has been proposed between the human Wa-like and porcine strains (16). Such an elegant whole genome genotyping system has not yet been established for non-RVA groups.

Determining the sequence of the 18.5kb segmented genome for rotaviruses by standard methods can be biased and cumbersome, requiring an initial PCR step to identify and select primers specific for the RV genogroup and/or strain, with the 11-segment genome providing an additional complication for primer design. Such primer-based sequencing strategies can be further complicated by reassortment possibilities not predicted by the initial PCR typing, leading to sequencing failure of atypical and un-typeable RV strains whose frequency can vary by location, season and environment (10, 17–21). Next-generation sequencing has been recently employed for whole-genome sequencing of RVA with initial 11 PCR amplifications (22–24); however, a single robust platform for whole-genome deep sequencing of multiple rotavirus genogroups without prior genotype information would be useful. Routine identification of circulating RVA can be performed using commercial enzyme immunoassay kits (based on inner capsid protein VP6) and RT-PCR diagnostic and genotyping assays (based on outer capsid proteins VP4 and VP7) (3, 25). In addtion, specific, rapid and cost-effective assays are lacking forthe detection of less common rotaviruses such as viruses in group B, C and H (RVB, RVC, RVH), hindering our understanding of molecular epidemiology of these viruses and challenging efforts of genomic sequencing, particularly in resource-limited countries (3, 25).

The rotavirus vaccines (Rotarix and RotaTeq) have been available since 2006(26, 27), and offer a variable degree of protective immunity against human RVA infections. Reduced RVA vaccine efficacy has been observed in resource-limited countries in comparison to developed countries (28–31). The mechanism responsible for reduced vaccine efficacy in these settings is unclear, but may in part be due to local circulation of genetically and antigenically divergent RVA or zoonotic strains in developing countries (10). Similar to other segmented viruses (32), genetic reassortment has been observed in RVs yielding significant genetic diversity, including a number of cross-species reassortants (3, 4, 25). Hence, assessment of all 11 genome segments through full virus genome sequencing is essential for monitoring the overall RV genomic diversity, complex evolutionary dynamics and the emergence of novel andzoonotic reassortants that may compromise vaccine protection (11, 14, 15).

Vietnam is a low to middle income country located in Southeast Asia and is considered one of the global hot spots of emerging infectious diseases (33). Diarrhoea is the fourth most common cause of mortality in children <5 years of age, accounting for 12% of deaths in this age group in 2013 (34). Among all diarrhoeal pathogens, RVA is responsible for 44% - 67.4% of all childhood diarrhoea cases requiring hospitalisation (35–48). Contrary to the clinical and public health importance, vaccination against RVA is currently not part of the Extended Program on Immunisation for Vietnamese infants. Additionally, diagnosis for rotaviruses in diarrhoeal cases in humans and animals is not routinely performed and systematically genomic surveillance of circulating human and animal rotaviruses is limited. This leads to relatively little data on the overall rotavirus prevalence and diversity in human and animal populations and their contribution to human infections and their potential to compromise RVA vaccine protection. Given the tropical climate of Vietnam prone to flooding, the frequent close human-animals living proximity and high prevalence of infectious diseases, we hypothesized that rotavirus zoonosis may occur in the region but is under-investigated and under-characterised. There is no report on the overall prevalence of other RVs (non-RVA) in both humans and animals in this region. To address this knowledge gap, we used focused sampling within human healthcare and animal farming populations, combined with high-throughput primer-independent direct genome sequencing from clinical materials (49) to document rotavirusdiversity and transmission within and between humans and animals in a region of Vietnam.

## MATERIALS AND METHODS

**Study setting and design.** Human and porcine faecal samples were collected from Dong Thap, a peri-urban province located in the south of Vietnam in the Mekong Delta region (see map, Supplementary Figure S1). The human subjects were diarrhoeal patients (N = 146) admitted to Dong Thap Provincial Hospital in the period from October 2012 to January 2014; a stool specimen was collected from each individual within 24 hours of hospital admission to avoid confounding by nosocomial infections. A total of 150 porcine faecal samples were randomly selected from a collection of porcine stool samples from pig farm baseline surveillance samples collected across the same province from January 2012 to April 2013. For 4 pigs in farms where no faecal specimens were obtained, a boot swab was collected (pig ID 12087_38, 14152_6, 14150_53 and 14250_12). All collected faecal samples were stored in aliquots at −80°C until further processing. Ethical approval for the study was obtained from the Oxford Tropical Research Ethics Committee (OxTREC Approval No. 15-12) (Oxford, United Kingdom), the institutional ethical review board of Dong Thap Provincial Hospital (DTPH) and the Sub-Department of Animal Health Dong Thap province (Dong Thap, Vietnam).

**Mapping of the patient residential and pig farm addresses.** The residential district centroid was recorded for enrolled human patients to maintain participant anonymity, while the exact geographical location was recorded for the pig farms using an eTrex Legend GPS device (Garmin, United Kingdom). The decimal degrees of latitude and longitude were enteredin a confidential database and kept separate from patient metadata so that patient identities could not be revealed based on the residence locations. These addresses were then validated in Google Earth Pro (https://www.google.com/earth/) and finally visualised in QGIS v2.2.0 (http://www.qgis.org/en/site/) overlaid with province-specific geographic data.

**Sample preparation and nucleic acid extraction.** Total nucleic acid extraction was performed as previously described (49–51). Briefly, 110 μL of a 50% stool suspension in PBS was centrifuged for 10 minutes at 10,000 X g. Non-encapsidated DNA in the samples was degraded by addition of 20 U TURBO DNase (Ambion). Virion-protected nucleic acid was subsequently extracted using the Boom method (52). Reverse transcription was performed using non-ribosomal random hexamers (53) that avoid transcription of rRNA, and second strand DNA synthesis was performed using 5U of Klenow fragment 3’-5’ exo^-^ (New England Biolabs). Final purification of extracted nucleic acids was performed with phenol/chloroform and ethanol precipitation.

**Library preparation and sequencing.** Standard Illumina libraries were prepared for each sample. In short, nucleic acids in each sample were sheared to 400-500 nt in length, each sample’s nucleic acid was separately indexed and samples were multiplexed at either 7 samples per MiSeq run or 96 samples per HiSeq 2500 run, generating 2-3 million 149 nt (MiSeq) or 250 nt (HiSeq) paired-end reads per sample.

***De novo* assembly and identification of viral genomes.** Raw sequencing reads were filtered to remove low quality reads (Phred score >35) and trimmed to remove residual sequencing adapters using QUASR (54). The reads were assembled into contigs using *de novo* assembly with SPAdes (55) combined with sSpace (56). RV-encoding contigs and other mammalian virus contigs were identified with a modified SLIM algorithm (49) combined with ublast (57). Coverage was determined for all contigs harvested to filter any process contamination sequences in each run, followed by additional filtering for minimum contig size cutoff (300 nt). Partial but overlapping contigs were joined into full-length sequences using Sequencher (Gene Codes Corporation, USA), and any ambiguities were resolved by consulting the original short reads. Final quality control of genomes included a comparison of the sequences, open reading frames (ORFs) and the encoded proteins with reference sequences retrieved from GenBank.

**Genotyping and phylogenetic reconstruction.** Assembled RVA sequences were genotyped using the online genotyping tool, RotaC v2.0 (http://rotac.regatools.be) (58), according to the guidelines for precise RVA classification using all 11 genomic segments (11). The resulting RVA, RVB, RVC and RVH sequences were combined with additional full-length or nearly full-length sequences from previous Vietnamese studies (if available) and global representatives retrieved from GenBank. The complete genomes from the vaccine components of the monovalent vaccine Rotarix (59) and the pentavalent vaccine RotaTeq (60) were retrieved from GenBank for phylogenetic reconstructions of all 11 RVA segments. Sequences were aligned using MUSCLE v3.8.31 (61) and manually checked in AliView (62); aligned sequences were trimmed to complete ORFs for subsequent analyses. Evolutionary model testing was implemented in IQ-TREE v3.10 (63) using the Akaike Information Criterion (AIC) to determine the best-fit models of nucleotide substitution for all genomic segments. Maximum likelihood (ML) phylogenetic trees were then inferred in IQ-TREE v3.10 with 500 bootstrap replicates under the best-fit model of evolution according to AIC (Supplementary Table 1 summarised the models determined for all segments). Resulting trees were visualised and edited using FigTree v1.4.2 (http://tree.bio.ed.ac.uk/software/figtree/). Genetic distances (p-uncorrected) were estimated using Geneious v9.0.4 (Biomatters Ltd).

**Bayesian analysis for RVA NSP3 genotype T7.** Available RVA sequences of T7 type (NSP3 segment) retrieved from GenBank and new sequences obtained in this study were aligned using MUSCLE v3.8.31 (61), manually checked in AliView (62), and trimmed to complete ORF. A maximum likelihood phylogenetic tree was constructed under the GTR+Γ_4_ model of substitution in IQ-TREE v3.10 (63). The molecular clock model was assessed in Path-O-Gen v1.4 (http://tree.bio.ed.ac.uk/software/pathogen/), assessing the linear regression between root-to-tip divergence and the date of sampling (year; as data on day and month were not available for GenBank sequences). A Bayesian Markov chain Monte Carlo (MCMC) approach was then performed in BEAST v1.8.0 (64) with three independent chains, using relaxed lognormal molecular clock under HKY85+Γ_4_ substitution model with a Bayesian SkyGid population process for 100 million generations chain with sampling performed every 10,000 runs. These triplicate runs were then combined using LogCombiner v1.8.0 (available within the BEAST package) with a removal of 10% burn-in, and analysed in Tracer v1.6 (http://tree.bio.ed.ac.uk/software/tracer/) to ensure all parameters had converged with effective sample size (ESS) values >200 and to estimate the mean evolutionary rates across branches. Maximum clade credibility trees were annotated using TreeAnnotator v1.8.0 (BEAST) and visualised in FigTree v1.4.2.

**Bayesian analysis for RVH VP6 gene.** A maximum likelihood phylogenetic tree was inferred for all available RVH VP6 sequences retrieved from GenBank (N=39) and from this study (N=5) under the GTR+Γ_4_ model of substitution in IQ-TREE (63). A time-scaled phylogeny was inferred for sequences within the porcine lineage. Highly similar sequences were removed before running Bayesian analyses (strains BR59, BR60, BR61, BR62, BR63, NC7_64_3, OK5_68_10). The molecular clock model was assessed in Path-O-Gen, and a Bayesian MCMC approach was then performed on the final set of sequences in BEAST (64), employing a relaxed lognormal molecular clock under HKY8+Γ_4_ substitution model with a non-parametric Guassian Markov Random Fields (GMRF) Bayesian Skyride population with tip dates defined as year, month, day of strain collection. Analyses were run in triplicate for 50 million generations with sampling performed every 5,000 generations. Triplicate runs were combined using LogCombiner with a removal of 10% burn-in, followed by analyses in Tracer, TreeAnnotator and FigTree as outlined in the aforementioned section.

**GenBank accession numbers.** All sequences generated in this study were deposited into GenBank under accession numbers KX362367 - KX363442. Illumina raw read sets are available at the European Nucleotide Archive under submission ERR471259 - ERR477293, ERR689707 - ERR767572, ERR775471 - ERR780002, ERR780013 - ERR780019, ERR956666, ERR956667, ERR962074, ERR1300950 - ERR1301100.

## RESULTS

### Overall diversity of rotaviruses in human and pigs

Sequencing of human enteric samples from acute diarrhoeal patients admitted to Dong Thap Provincial Hospital from 2012-2014 yielded 60 *de novo* assembled RVA genome sequences from 146 samples (41.1%). No other RV genogroups were found in these human stool samples (Table 1). The same methods applied to 150 porcine faecal samples collected within the same geographic region (Supplementary Figure 1) identified 31 rotaviruses from 4 different RV groups (A, B, C and H) in a total of 150 samples (20.7%). These *de novo* assembled sequences included 13 RVA (41.9%), 7 RVB (22.6%), 6 RVC (19.4%) and 5 RVH (16.1%) (Table 1). The length of each assembled sequence was determined and expressed as percentage length coverage (length of assembled sequence divided by expected full length of that segment) for the corresponding segment (Figure 1). In samples where 2 distinct contigs were assembled for a segment (e.g. mixed infections), only the longer assembled contig was reported in the heatmap of segment coverage for the purpose of clarity (Figure 1). The overall length coverage in human RVA sequences was higher than porcine RVA, RVB, RVC and RVH, possibly be due to differential viral load or sample quality.

**Figure 1.**
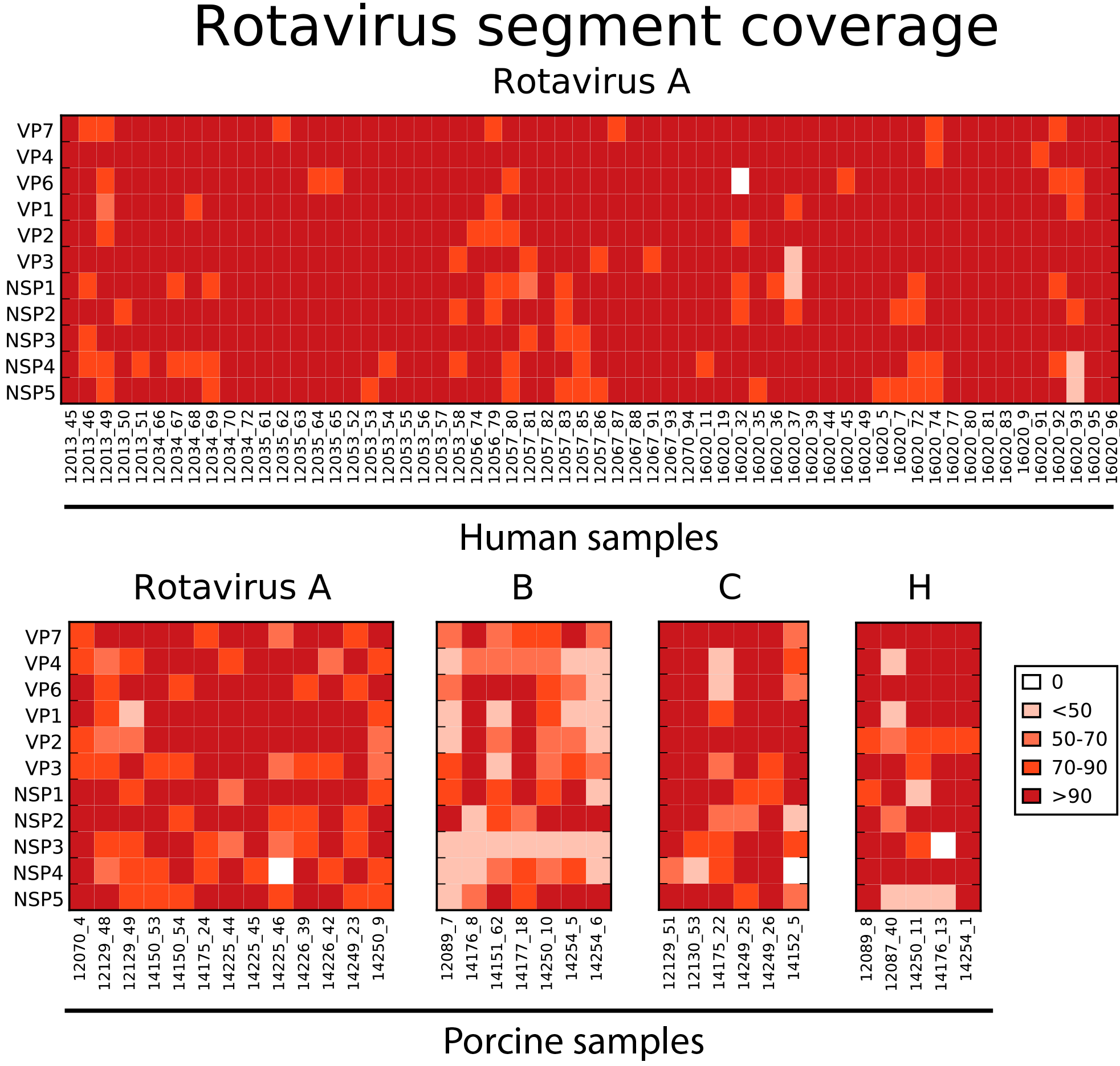
Heat map of rotavirus sequence length coverage by segment detected in all
samples. The sequence length coverage for each segment of all assembled rotaviruses by deep sequencing was calculated and expressed as [(length of assembled contig in nt)/(full-length of that segment in nt)]×100. Colour code for %genome coverage is indicated in figure key ranging from low (pale orange) to high (dark red). The value of 0 indicates that contig sequence for that segment was not identified or did not pass the stringent quality control criteria including % reads mapped and contig length (see Materials and Methods). All RVA sequences detected in human samples were shown in the first panel, with each column representing a sample and each row showing each RV segment. Similarly, porcine rotavirus samples wereshown, each panel representing RVA,RVB, RVC and RVH sequences with the segment names given vertically and sample IDs horizontally.

**Table 1.** Number of samples tested and number of samples yielded up to full or nearly fullRV genomic sequences.

| Host | RVA | RVB | RVC | RVH | Total samples tested |
| --- | --- | --- | --- | --- | --- |
| Human | 60<br>(41.1%) | 0 | 0 | 0 | 146 |
| Pig | 13<br>(8.7%) | 7<br>(4.7%) | 6<br>(4%) | 5<br>(3.3%) | 150 |

Host indicates human or pig host from which the faecal samples were collected. Percentages of full or nearly full RV genomes were given in brackets. RVA: rotavirus genogroup A; RVB: group B; group C; RVH: group H.

### Human and porcine RVA genotype constellations

The genotype constellations were determined for all RVA strains according to established guidelines from the Rotavirus Classification Working Group (11, 58). Among the human RVA, the two most common constellations were G1-P[8]-I1-R1-C1-M1-A1-N1-T1-E1-H1 (N = 33; Wa-like constellation) and G2-P[4]-I2-R2-C2-M2-A2-N2-T2-E2-H2 (N = 12; DS-1-like constellation) (Figure 2). Reassorted RVA strains between Wa-like and DS-1-like constellations were found in 4 human diarrhoeal patients, including G1-P[8]-I2-R2-C2-M2-A2-N2-T2-E2-H2 (N = 3) and G2-P[8]-I2-R2-C2-M2-A2-N2-T2-E2-H2 (N = 1). Interestingly, the genotype constellation G4-P[6]-I1-R1-C1-M1-A8-N1-T7-E1-H1 was found in 1 human and 1 pig sample. The closely related genotype constellation G4-P[6]-I1-R1-C1-M1-A8-N1-T1-E1-H1, which differs only by the NSP3 segment, predominated among the porcine RVA mono-infections (N = 6) (Figure 2). Among all porcine RVA strains, the internal core gene cassette of R1-C1-M1-A8-N1-T1-E1-H1 (representing genotype of VP1-VP2-VP3-NSP1-NSP2-NSP3-NSP4-NSP5) was relatively conserved, withthe exception of co-circulation of T1 and T7 genotypes in the NSP3 segment. The genotypes of capsid proteins (VP7-VP4-VP6) were more diverse in pigs, including G4-P[6]-I1 (6 strains combined with NSP3 T1 and 1 strain with NSP3 T7), G5-P[13]-I5 (N = 2),G9-P[23]-I5 (N = 1) and G11-P[23]-I5 (N = 1).

**Figure 2.**
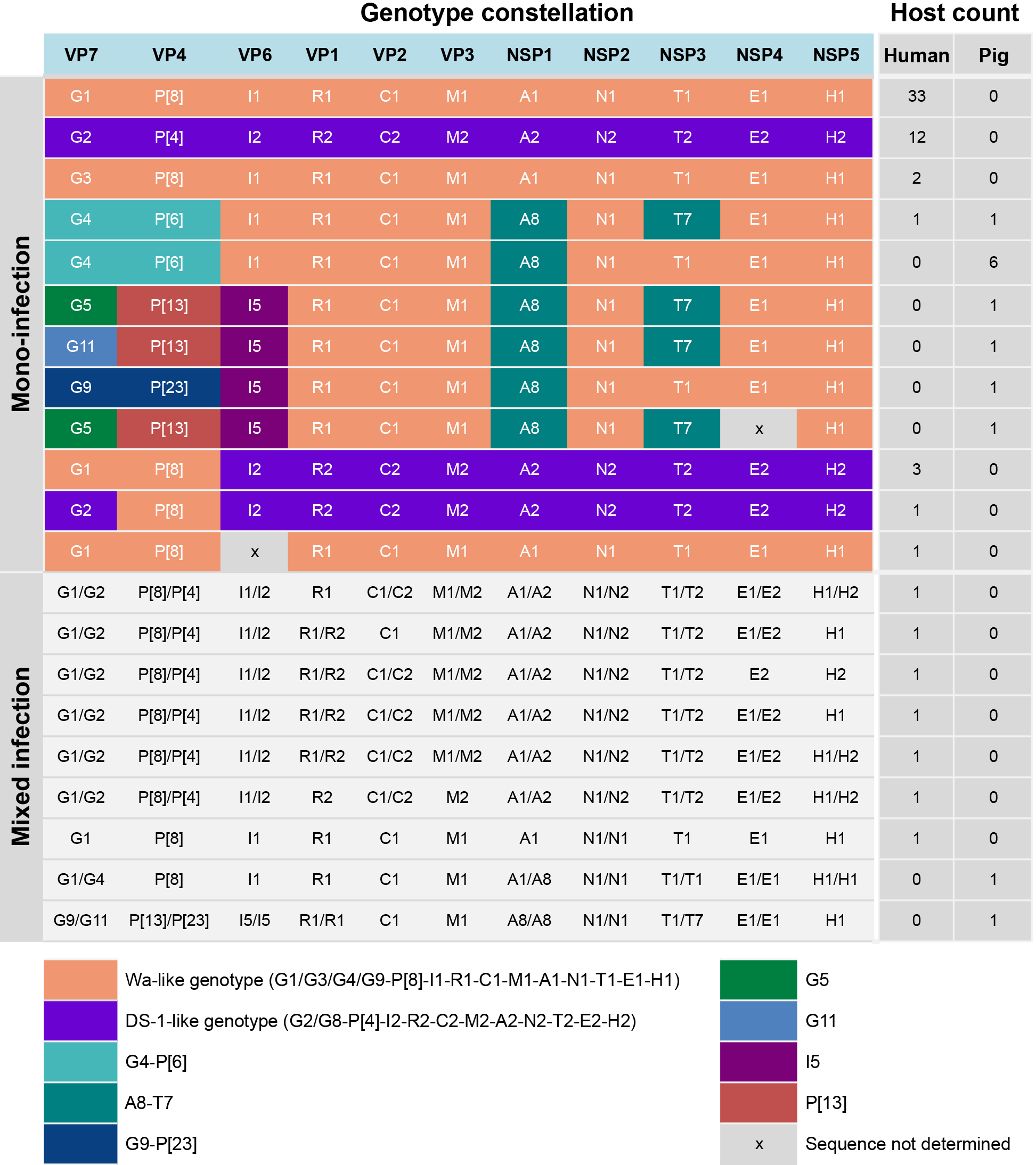
Genotype constellations of assembled RVA genomes. The genotypes ofall assembled sequences were determined according to the guidelines of Rotavirus Classification Working Group, representing the genotype of VP7-VP4-VP6-VP1-VP2-VP3-NSP1-NSP2-NSP3-NSP4-NSP5.For mono-infection, each row represents one genotype constellation with the colour block used to illustrate different genotype patterns, such as common human types of Wa-like (orange) and DS-1-like (purple), and other less common genotypes shown in other colour blocks asindicated. The number of each of the genotype constellation identified in samples from humanand pigs were given in the column Host count for Human and Pigs,respectively. For mixed infection where 2 distinct contigs were assembled for at least one segment, the genotypes of both contigs were given and indicated the host where strains were
identified.

Mixed infections were identified in 9 samples, 7 in humans and 2 in pigs (Figure 2), with mixed infection being defined as the detection of two assembled but genetically distinct contigs in at least one segment with sufficient contig coverage to exclude potential process contamination among samples in the same run. The two homologous contig segments identified in mixed infections can have different or the same genotype; for example a mixed infection reported in an individual pig (sample ID 12070_4) contained 2 homologous VP7 segments, NSP1, NSP2, NSP3, NSP4 and NSP5 bearing the constellation of G1/G4-P[8]-I1-R1-C1-M1-A1/A8-N1/N1-T1/T1-E1/E1-H1/H1 (Figure 2 and Supplementary Figure S2). Another porcine sample was found with a mixture of G9/G11-P[13]/P[23]-I5/I5-R1/R1-C1-M1-A8/A8-N1/N1-T1/T1-E1/E1-H1 (sample ID 14150_53); however, it is important to note that this particular sample was a boot swab of faecal material in a cage-type pigsty, thus there is the possibility that the sample represents mixed environmental virus from more than one pig. Mixed human RVA infections typically contained genotype 1 and 2 (Wa-like and DS-1 like, respectively) viruses.

### Phylogenetic diversity of local human and porcine RVA

Phylogenetic trees were inferred for each RVA segment (Figure 3A-B and Supplementary Figure S3A-B) from assembled sequences in this study along with full-length sequences from previous studies in Vietnam, reference sequences in GenBank and sequences from the RVA vaccine formulations (RotaRix and RotaTeq). The local sequences clustered primarily by genotype as expected (Figure 3A-B and Supplementary Figure S3A-B); for example, VP7 G1 sequences in this study clustered with other G1 sequences from other regions and our G2 sequences clustered with other G2 (Figure 3A). Sequences within the G4 genotype fell into 2 sub-lineages, with the human strain (16020_7) and porcine strains from this study clustering into one common sublineage (Figure 3A). The mixed infection in the pig (12070_4) described above comprised 2 distinct contigs for the VP7 segment (belonging to the G1 and G4 genotypes), with the G1 sequence clustering within the human G1 lineage and the G4 sequence falling into a lineage with other G4 porcine sequences from this study (Figure 3A). Similar observations were seen in the phylogenetic tree for VP4 sequences (Figure 3B) and other gene segments (Supplementary Figure S3A-B). It is also noteworthy that sequences from the vaccine strains (Rotarix and RotaTeq) were relatively distinct from the Vietnamese RVA sequences reported here, particularly for genotypes G5, G9, G11, P[6], P[13], P[23] of the two neutralizing antigens, VP7 and VP4 (Figure 3A-B). Comparison of the amino acid sequences of VP7 and VP4 of the local strains to the Rotarix and RotaTeq vaccine sequences indicated a number of amino acid differences observed across the length of the proteins and particularly in the antigenic epitopes of VP7 and VP4 (Supplementary Figure S7). Taken together, multiple RVA genotypes co-circulate in human and pigs in this location; many of these genotypes are genetically dissimilar to currently used vaccine components.

**Figure 3.**
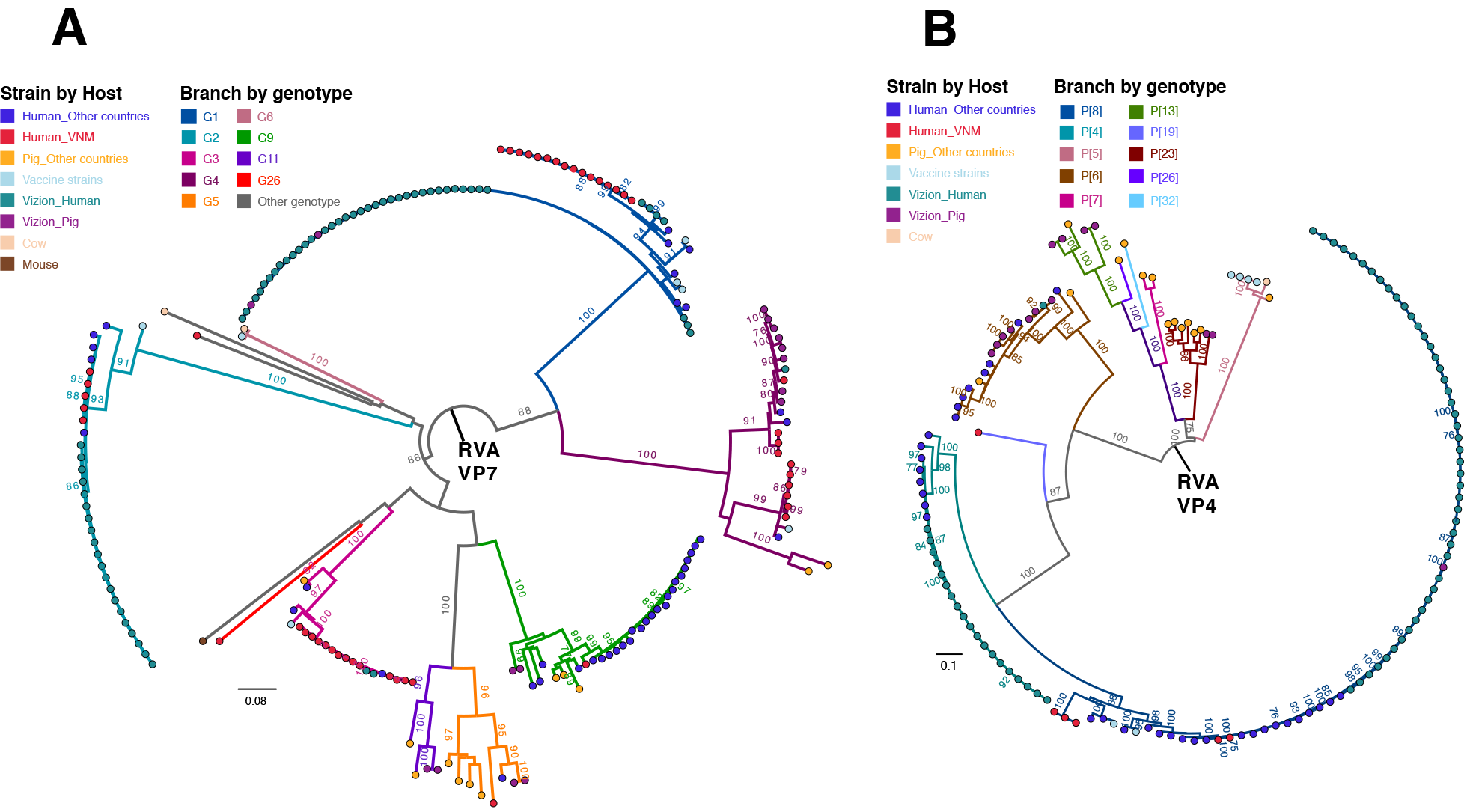
Maximum-likelihood phylogenetic trees inferred from the assembled nucleotide
sequences for RVA VP7 and VP4 genes. **(A)** Maximum-likelihood tree of VP7 gene showed genetic relationships between sequences from this study and additional sequences of corresponding segments from GenBank. Branches and strain names were coloured according to genotype and host, respectively. Tree is mid-point rooted for the purpose of clarity and bootstrap values of ≥75% are shown for major nodes only. All horizontal branch lengths aredrawn to the scale of nucleotide substitutions per site.**(B)** Maximum-likelihood tree of VP4 gene showed genetic relationships between sequences from this study and additional sequencesof corresponding segments from GenBank. The pattern of tree visualisationis consistent withVP7 tree, see description of Figure 3A for more information

### Putative zoonotic infection of human with a porcine-human RVA virus

An atypical RVA genotype constellation of G4-P[6]-I1-R1-C1-M1-A8-N1-T7-E1-H1 was found in both a human patient (16020_7) and a weaning pig (14250_9), whose geographical distance (between the residence and the farm) were approximately 35km apart (Figure 4A-B). The genotype constellation of the core gene cassette (R1-C1-M1-A8-N1-T7-E1-H1) was also identified in 4 pig samples collected from 2 other farms (Figure 4A); the farms that raised these pigs are also about 35km away from the residential location of the 16020_7 case (Figure 4B). RVA strains with the aforementioned genome constellation have been identified in paediatric diarrhoeal patients in paediatric diarrhoeal patients in Hungary (65), Argentina (66), Paraguay (67), and Nicaragua (68), and were similarly thought to be of zoonotic origin (porcine-like). Given the diversity of geographic locations of reported zoonotic cases over a 15-year period, it is difficult to determine if this strain is sustained as a rare variant in human-to-human infections or has undergone multiple cross-species jumps from a porcine reservoir or an intermediate host.

**Figure 4.**
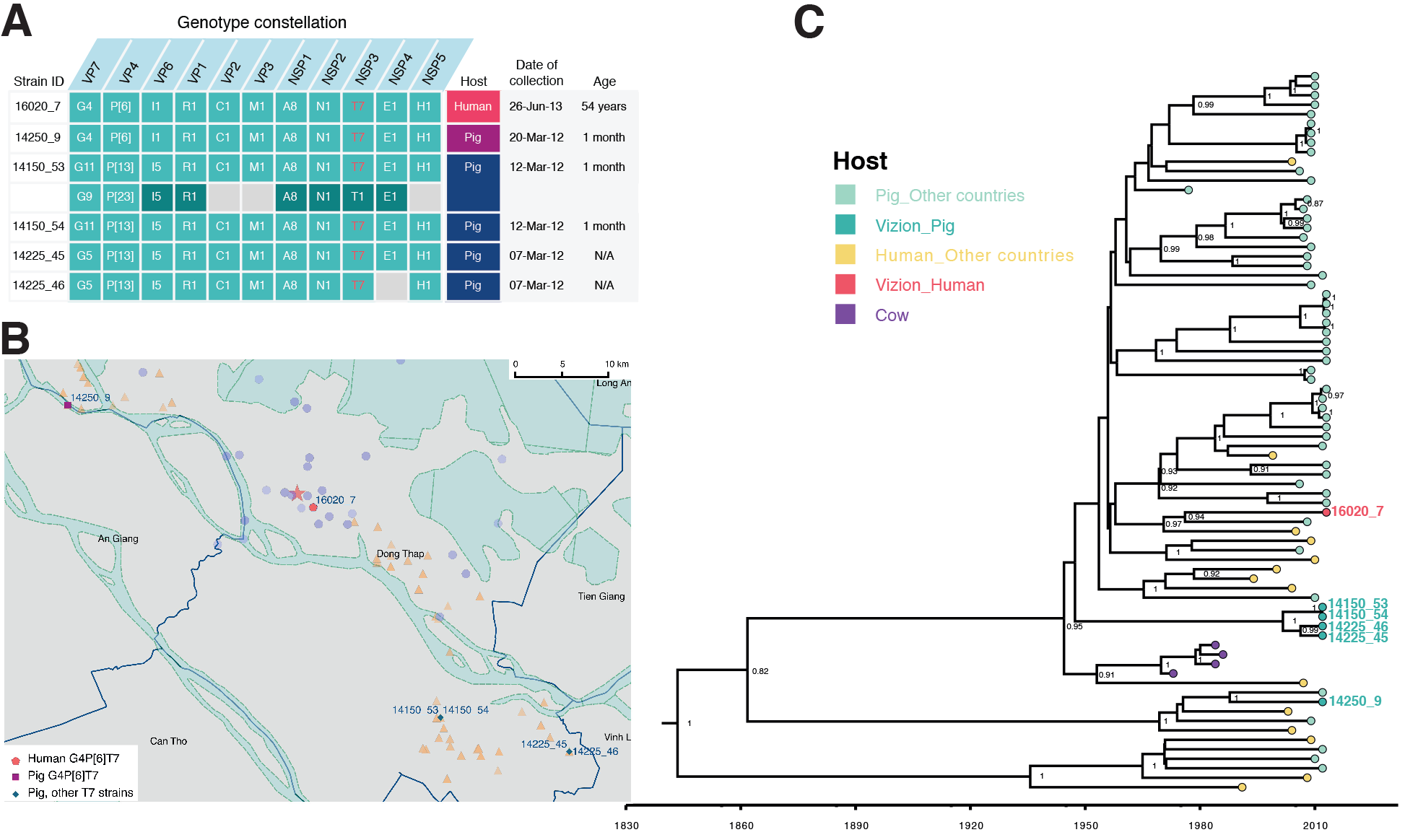
The analysis of RVA zoonotic strain G4P[6]T7 in the study. **(A)**Thegenotype constellation of the case in investigation 16020_7 and other porcine RVA strains with the NSP3 T7 genotype. Details on dates of collection and age of host are given. The colour coded in the host column is consistent for colour illustration of corresponding case in the map (panel B) in this figure. **(B)** The geographical location of the human case’s residency and pig farms that raised the pigs infected with RVA NSP3 T7 strains overlaid on the total sampling area (as shown in Supplementary Figure S1). The colouring of the human case and pig farms is consistent with colour code presented in the column “Host” in panel of this figure. The red star indicates the Dong Thap Provincial Hospital where diarrhoeal patients were admitted. The map scale bar is shown in the units of geometrickm. See Supplementary Figure S1 for more information. **(C)** The time-resolved phylogenetic treeof RVA NSP3 T7 genotype sequences comparing local versus global sequences. Reference sequences were retrieved from GenBank (N=69, excluded the duplicate sequence for TM-a strain (JX290174) and BP1901 (KF835960) which is 15 aa shorter than the complete ORF). Strains were coloured according to the host species from which the strain was identified. Sequences identified from this study were highlighted in blue (human case 16020_7) or in yellow (porcine sequences). The first T7 sequence identified (in a cow in 1973) was indicated in purple. Posterior probabilities of internal nodes with values ≥0.75 are shown and the scale axis indicates time in year of strain identification.

Phylogenetically, all 11 segments of the human 16020_7 and porcine 14250_9 viruses belonged to lineages comprising porcine and/or porcine-origin human sequences (Figure 3A-B and Supplementary Figure S3A-B). Genetic distance suggested that the strain 16020_7 was most similar to porcine strains: TM-a (for VP1; 96% nt similarity), CMP45 (NSP3; 93%), and porcine-origin human strain 30378 (NSP2; 99%) (Figure 5). The remaining segments were most similar to porcine RVA strains obtained from this study, including the porcine 14150_53 (NSP1; 98.7% and NSP4; 99.1%) and 14225_44 (VP2; 99.2% and NSP5; 99.1%) strains. The capsid proteins of 16020_7 were most similar to the VP7 sequences of porcine samples 12129_48, 12129_49 and 12070_4 (G4 type; 97.2%), to the VP4 of pig 14226_39 (98.7%) and VP6 of pig 14226_42 (99.5%) (Figure 5). The porcine sample 14250_9, despite possessing the same genotype constellation as 16020_7, shared the highest nucleotide homology in only 2 internal genes, VP3 (97.5%) and NSP5 (99.1%), to the corresponding segments of strain 16020_7. Compared to the RVA vaccine Rotarix, 16020_7 was relatively dissimilar, sharing as low as 75.1% and 75.6% nt similarity for the VP4 and VP7 segments, respectively (Figure 5).

**Figure 5.**
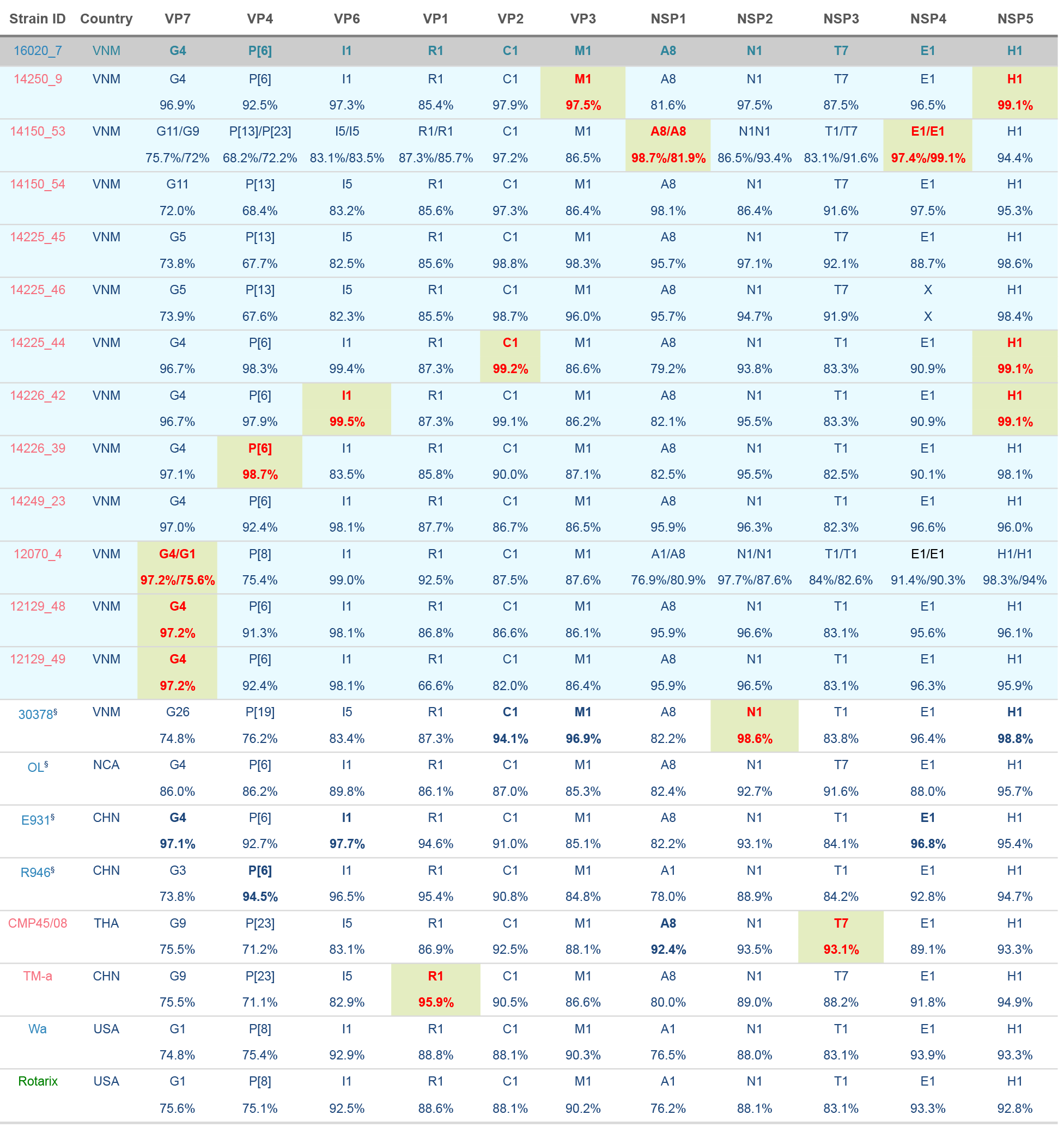
Genotype constellations of RVA 16020_7 compared with representative human and animal RVA of known genotypes. Segments are bold in red and shaded in green to indicate the segments with highest nucleotide sequence similarities to that of strain 16020_7.The strain was coloured according the host from which strain was identified, blue indicateshuman host,pink for pigs and green for vaccine component. All porcine samples from this study were shaded in light blue and the human case of interest (16020_7) was shaded in grey. §Human strains were previously shown to have porcine origin. Country of isolation abbreviation,
 VNM: Vietnam; NCA: Nicaragua; CHN: China; THA: Thailand; USA: United States of America.

In this unusual genotype constellation reported, the NSP3 T7 type is a rare genotype that was first identified in a cow in Great Britain in 1973 (69), then in a bovine-like human strain (70), and later in pigs (71), porcine-bovine human reassortant (72) and porcine-like human strains (65–68, 73) in various geographical locations. The inferred evolutionary rate of RVA NSP3 sequences bearing T7 genotype was 1.3261 × 10^−3^ substitutions per site per year (95% highest posterior density (HPD): 8.624 × 10^−4^ − 1.793 × 10^−3^), which is slightly lower than the estimated evolutionary rates for RVA VP7 capsid gene of 1.66 × 10^−3^ and 1.87 × 10^−3^ substitutions/site/year for G12 and G9 genotypes, respectively (74). The time-stamped MCC tree also indicate an inter-connection among different host species indicating several host jump events particularly between pig and human hosts, suggesting that viral zoonotic chatter may occur more frequently than hitherto reported (Figure 4C).

### Rotavirus group H (RVH), group B (RVB) and C (RVC)

RVH was identified in five Vietnamese pigs (3.33%; 5/150) at several time points and locations with no temporal or geographical associations (Figure 6), suggesting that these infections were sporadic and not linked to a single local outbreak. Furthermore, phylogenetic trees were inferred for all RVH segments to investigate the genetic diversity, comparing the RVH strains identified in this study with RVH sequences retrieved from GenBank (Figure 6 and Supplementary Figure S6). In general, RVH sequences typically clustered according to the host species, i.e. all porcine RVH sequences belonged to a lineage that is separated from human or cow RVH lineages. Within the porcine clade of the VP6 gene (Figure 6), sequences fell into two lineages: one lineage comprising sequences from USA and Japan, and the other lineage of Brazilian and Vietnamese sequences. The evolutionary rate was estimated to be 5.195 × 10^−3^ substitutions per site per year for sequences in the porcine lineage of RVH VP6 (95% HPD: 1.865 × 10^−3^ - 8.976 × 10^−3^).

**Figure 6.**
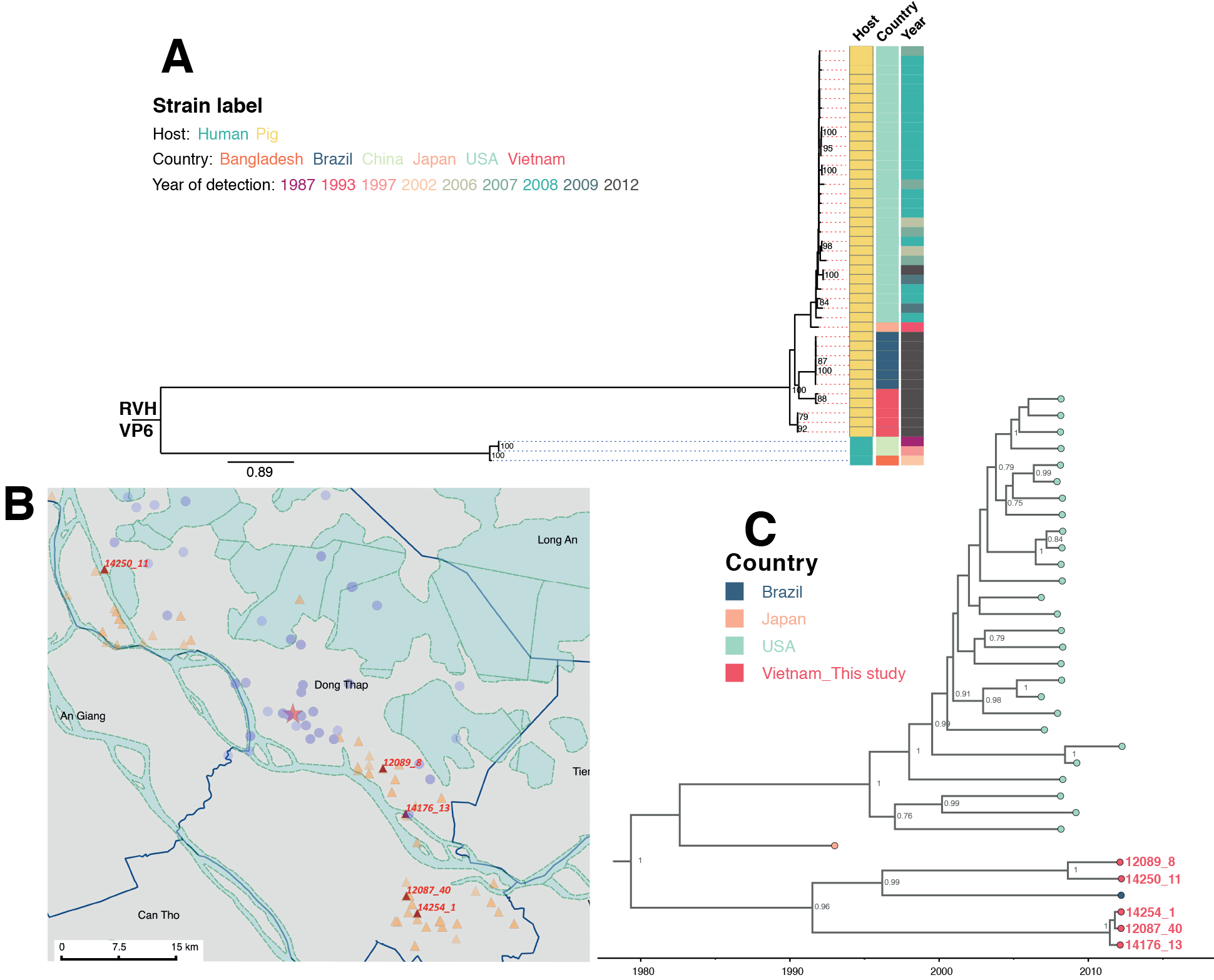
The analysis of RVH VP6 gene segment. **(A)** Maximum-likelihood phylogenetic tree of RVH VP6 sequences. The RVH VP6 sequences in this study (N=5) were compared with available full-length RVH VP6 sequences retrieved from GenBank (N=39). Tree is mid-pointrooted for the purpose of clarity and only bootstrap values of ≥75% are shown. All horizontal branch lengths are drawn to the scale of nucleotide substitutions per site in the tree. Strains were colour coded according to the host associated with the strain, the country where the strains were identified and the year of strain detection. **(B)** The geographical locationsof pig farms that raised the pigs infected with RVHidentified in this study. The farms were illustrated as red triangles with strain ID given, overlaid on the overall sampling area as shown in Supplementary Figure S1. The red star indicates the Dong Thap Provincial Hospital. The map scale bar is shown in the units of geometric km. Refer to Supplementary Figure S1 for more information on the background and provincial features colouring. **(C)** Time-resolved phylogenetic tree of porcine RVH VP6 sequences comparing local versus global sequences. Strains were coloured by the country where strains were identified. Porcine sequences identified from this study were highlighted in red, with strain ID given to link with geographical locations on map in panel C of this figure. Strains from Brazil were coloured in dark blue; orange indicates the Japanese porcine strain, and light green refers to the porcine strain from the US. Posterior probabilities of internal nodes with values ≥0.75 are shown and the scale axis indicates time in year of strain identification.

RVB was found in 7 pigs (4.67%; 7/150) and RVC was identified in 6 pigs (4%; 6/150). Phylogenetic trees of all segments of RVB and RVC showed that the local porcine sequences belonged to lineages comprising porcine sequences from other geographical locations for RVB (Figure 7 and Supplementary Figure S4) and RVC (Figure 7 and Supplementary Figure S5). In both RVB and RVC groups, the porcine lineages were relatively distant from lineages comprising of human sequences (Figure 7 and Supplementary Figure S4 and S5).

**Figure 7.**
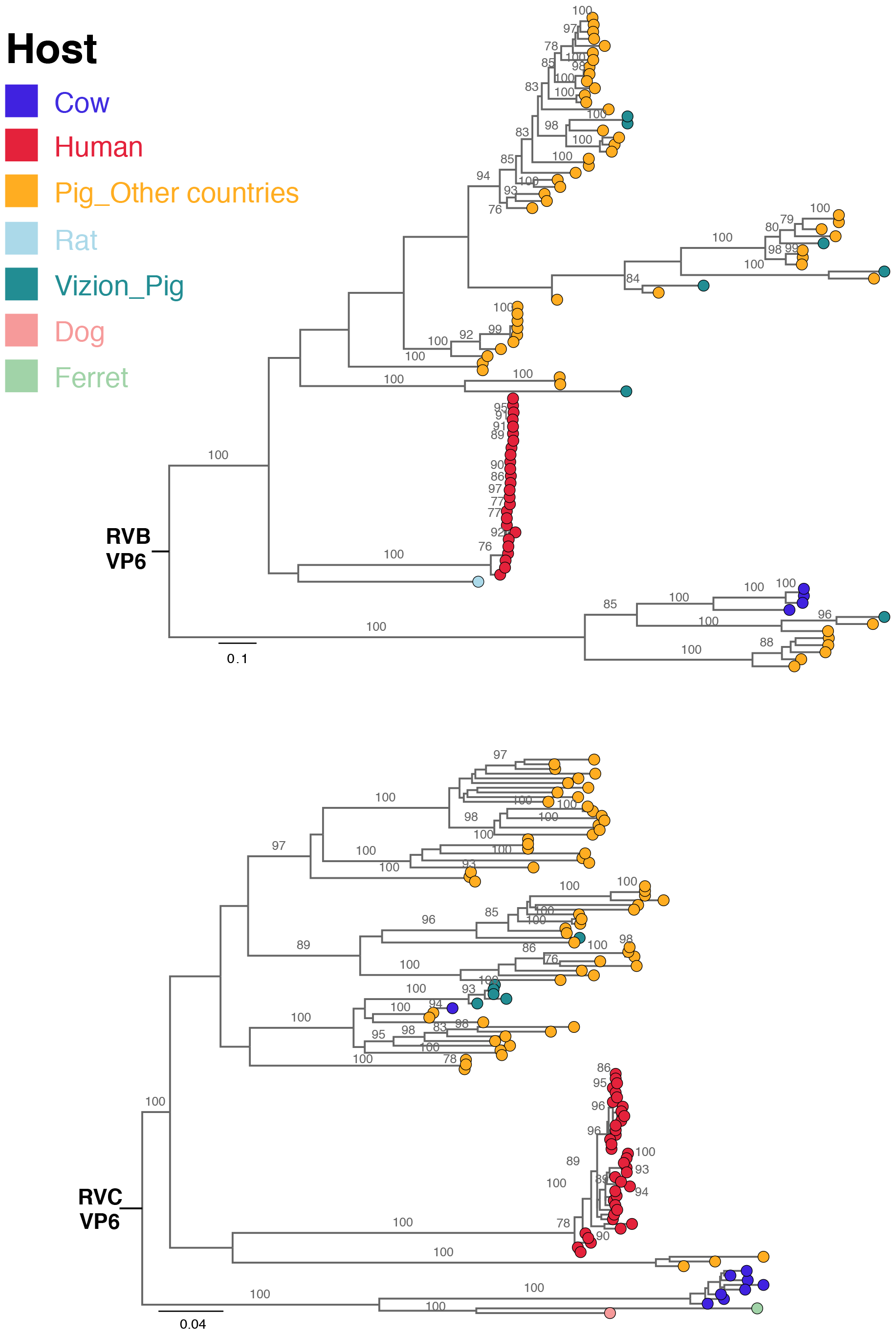
Maximum-likelihood phylogenetic trees inferred from the assembled nucleotide sequences for VP6 gene of RVB and RVC. Maximum-likelihood trees of RVB and RVC VP6 segments showed genetic relationships between assembled sequences from this study and full length reference sequences of corresponding segments retrieved from GenBank. Trees are mid point rooted for the purpose of clarity and only bootstrap values of ≥75% are shown. Scale barsare in the unit of nucleotide substitutions per site. Strains were coloured according to the hostspecies that the sequences were identified from

## DISCUSSION

This study represents the first unbiased genome-wide surveillance, targeting simultaneously multiple groups of rotaviruses infecting humans and animals in the same geographical location. Prior to this study, there were only 3 subgenomic RVC sequences (<300nt) and 9 complete or nearly complete RVA genomes reported from Vietnam in GenBank with no data on RVB and RVH. Data from the current study document genomic sequences from 60 human RVA and 31 porcine RV (group A, B, C and H), providing the largest available collection of genome sequences from human and pigs from a single location in general and from Vietnam in particular. This is also the first report and the first genome characterisation of RVB, RVC and RVH from Vietnamese pigs.

Among the RVA, we identified a human (sample 16020_7) and a porcine sample (sample 14250_9) with atypical RVA genotype constellation G4-P[6]-I1-R1-C1-M1-A8-N1-T7-E1-H1, detected for the first time in Vietnam and in Asia. This variant may have originated from a direct zoonotic transmission or from reassortment event(s) involving porcine and porcine-origin human strains. The human RVA strain (16020_7) was identified in a sample from a 54 year-old patient, admitted to the hospital due to acute diarrhoea. Rotavirus was the sole enteric pathogen identified from the stool sample and no other common viral and bacterial diarrhoeal pathogens were found by diagnostic testing (39, 75) for norovirus, astrovirus, sapovirus, adenovirus F and aichi virus, *Shigella spp., Salmonella spp.* and *Campylobacter spp.* (data not shown). Although adults can be infected with RVA, such infections in immuno-competent individuals are typically asymptomatic, self-limiting or cause mild disease (76). The rotavirus infection in this particular case required hospitalisation suggesting a moderate-severe end of the clinical spectrum of diarrhoeal disease. Although further studies are required to determine their significance and relevance in human and animal diseases, it is tempting to suggest that this atypical strain may be the cause of the moderate-severe diarrhoeal disease. The close proximity between humans and pigs and common use of river water (Mekong Delta River, Figure 4B) for daily activities and farming might present an enhanced risk of transmitting water-borne infectious pathogens, in this case providing a plausible zoonotic route of atypical rotavirus transmission.

Compared to RVA, human and animal infections with RVB, RVC and RVH are not well understood and the detection rates for these groups of viruses are relatively low (77). This is probably because the majority of rotavirus investigations have been focused on RVA given its clinical and public health relevance and the large genetic distance among these groups of rotaviruses as compared to RVA, which would likely be missed by commonly used diagnostic assays. Recently, there have been increasing numbers of reports on rotavirus groups B, C and H in animals (78–86) and humans (87–89), which possibly reflect improved molecular methods to detect these viruses rather than an actual increase in their prevalence. In Vietnam, the frequencies and relative role in human and animal disease of RVB, RVC and RVH viruses are not yet known. Although no human RVB, RVC and RVH were found in this study, the zoonotic potential of these rotaviruses groups cannot be ignored.

Our documentation of local RV genetic diversity and the potential of RV for zoonosis are highly relevant for the introduction of RVA vaccines into the region. The VP7 surface glycoprotein and VP4 spike protein are considered to be major antigenic targets for the protective host immune responses induced by RVA vaccines (73, 90, 91). Importantly, several amino acid changes were observed in defined antigenic sites of VP7 and VP4 proteins encoded by the local Vietnamese strains compared to available vaccines. Although a genetic comparison alone cannot predict if circulating Vietnamese viruses will be recognized and blocked or if the changes will allow infection in vaccinated subjects, the number of differences in antigenic regions suggests a compromised protective function of the vaccines in Vietnamese children. These data should be considered in light of the clinical observations of reduced vaccine efficacy (28–30). Future vaccine efforts may benefit from this increased knowledge of rotavirus diversity provided by this work.

Our study does have limitations. Firstly, the sample size of 146 human and 150 porcine samples is relatively small. Despite this size, we were able to identify a potential zoonotic infection. This provides a baseline frequency for zoonotic infections and suggest that this may be occurring at a higher rate than previously considered. Secondly, the disease status of the sampled pigs was not well defined and there was no follow-up beyond the sampling time point so the clinical spectrum of diarrhoeal disease (eg. mild, moderate or severe) in the pigs is unknown. However, the primary objective of this study was characterization of rotavirus pathogenesis or causation of the diarrhoeal disease. The presence of rotavirus material at sufficiently high titres to allow full genome sequencing is consistent with these animals being a common source of the virus for movement to other species. Our findings indicate that porcine faecal material is a source of novel and possibly zoonotic viruses.

It is likely that with the ubiquity and falling costs of sequencing, the unbiased virus sequencing described here will become an important component of infectious disease surveillance and rapid responses to outbreaks (92). The ideal sampling rate, sample numbers and geographical relationship between humans and animals for genetic surveillance are still being defined but the current work provides a good starting point for future efforts. Even within the relatively modest sample set of 296 human and animal enteric samples, a considerable RV genetic diversity was observed including a potential zoonosis. The integration of targeted sampling, sequencing and phylogeography or phylogenetics in different places in the world, perhaps informed by other risk mapping (33, 93) has the ability to inform surveillance and to monitor zoonotic pathogens in human and animals.

## Acknowledgements

We are grateful to all the patients and their families and the farmers in Dong Thap (Vietnam) that have participated in the study. We thank the Illumina C team at the Wellcome Trust Sanger Institute (Hinxton, Cambridge, United Kingdom) for their help in deep sequencing and Simon Watson and Pinky Langat (Wellcome Trust Sanger Institute) for technical assistance on running BEAST analyses. The complete list of the VIZIONS Consortium can be found at (94).

